# Arithmetic learning is associated with developmental increases in similarity between brain activity and artificial neural networks

**DOI:** 10.1101/2025.05.14.653407

**Authors:** Tomoya Nakai, Jérôme Prado

## Abstract

The ability to learn simple arithmetic is often thought to rely on the associative nature of human memory, where repeated exposure to arithmetic problems strengthens the relations between operands and outcomes. This view is reminiscent of learning in large language artificial neural networks (ANNs), where training involves statistical associations between symbolic inputs and outputs. Based on this parallel, we hypothesized that brain activity during arithmetic problem-solving should be increasingly similar to ANN-derived representations with learning and development. We used fMRI to investigate the relation between brain activity during single-digit addition problem-solving and ANN features in 104 participants across four age groups (8-, 11-, 14-year-olds, and adults). Vertex-wise encoding models were used to predict brain activity using latent features extracted from pre-trained ANNs. Results revealed that the ANN-based model better predicted arithmetic-related activity in older than younger participants in a specific region of the left precentral sulcus. Prediction accuracy was higher for smaller than larger problems, consistent with the idea that associations between problems may decrease in strength with problem size. Furthermore, ANN prediction patterns aligned with a model in which each addition problem was increasingly represented separately with age. On the one hand, these findings may support the idea that, with learning and development, some neural representations of arithmetic problems become discretely organized in memory and increasingly resemble ANN-like processing. On the other hand, the lack of relation between the ANN-based model and activity in other brain regions suggests some degree of dissimilarity between arithmetic processing in humans and ANNs, suggesting the existence of additional mechanisms for simple arithmetic learning in humans.

## Introduction

Our capacity to learn is largely thought to result from our ability to form and strengthen associations between elements. The idea that human memory is associative in nature has profound historical roots in both philosophy and psychology (Mandelbaum, 2015), supporting cognitive models that posit the existence of networks of associations in the human mind (Anderson & Bower, 2014). These accounts do not only assume that associative bonds can form between perceived elements, but also that bonds can form between elements and the outcomes of mental computations that are performed on them. The domain of simple arithmetic learning provides a compelling illustration of this principle. For example, it has been argued that there is a shift in the type of overt strategy children use to solve simple arithmetic problems, such as 5 + 2, with learning and experience (Ashcraft, 1992; Geary, Hoard, & Nugent, 2012; Lemaire & Siegler, 1995; Thevenot, Barrouillet, Castel, & Uittenhove, 2016). Up until the age of 9 or 10, children tend to solve these problems by using counting strategies, first with manipulatives (e.g., fingers) and then without (Wu et al., 2008). As children gain experience with these problems, overt counting strategies become less frequent and answers to these problems are progressively given quickly without any conscious report of a specific strategy (Ashcraft, 1982; Groen & Parkman, 1972). This has been traditionally interpreted as evidence of direct recollection from long term-memory, which would be made possible by the progressive forming and strengthening of associations between operands (e.g., 5 + 2) and outcomes (e.g., 7) over the course of multiple exposures during learning (Ashcraft, 1992; Chen & Campbell, 2018; Logan, 1988). In sum, mental arithmetic is a perfect example of a domain in which the associative nature of human memory is thought to support the shift from slow and relatively inefficient computations to fast and efficient retrieval of associations from memory.

In some ways, the hypothesized process of strengthening associations between operands and answers when children learn simple arithmetic is reminiscent of the way learning is achieved in large-language artificial neural networks (ANNs). ANNs are computational models inspired by biological neural networks (BNNs), usually organized in a directed network that may include multiple layers of neuronal units (Bishop, 2006; Goodfellow, I., Bengio, Y., Courville, A., & Bengio, Y., 2016). Each layer aggregates the outputs of preceding units, applies (typically nonlinear) transformation, and transmits the result to the next layer, enabling the network to learn complex input-output relationships. Much like natural language, mathematics involves processing sequences of symbols (Matsumoto & Nakai, 2023). Building on the recently developed Transformer model that can handle long-distance dependency in natural languages (Vaswani et al., 2017), several models have now been proposed to process mathematical expressions (Azerbayev et al., 2023; Peng, Yuan, Gao, & Tang, 2021; Schlag et al., 2019; Yang et al., 2024; Zong & Krishnamachari, 2023). Trained on large text corpora that include mathematical content, such ANNs output symbols based on statistical associations with the input symbol sequence (i.e., a math expression). Although ANNs can capture structural information in math expressions (Russin et al., 2021), their math ability may partly depend on the memorization of training samples (Yuan, Yuan, Tan, Wang, & Huang, 2023; H. Zhang et al., 2024). Therefore, ANNs process math expressions in a way that can be compared to the way older children and adults are often thought to solve simple arithmetic problems, i.e., by retrieving associations between elements.

Recent neuroimaging studies have observed similarities between the adult human brain and ANNs. For example, there is a correspondence between the hierarchy of human visual pathways and that of ANN layers in processing natural images (Cichy, Khosla, Pantazis, Torralba, & Oliva, 2016; Güçlü & van Gerven, 2015; Horikawa & Kamitani, 2017), where shallower layers correspond to low-level features (e.g., local contrast, edge, etc.) and deeper layers correspond to more complex information (e.g., entire shape). Similar hierarchical correspondence has been reported for auditory (Kell, Yamins, Shook, Norman-Haignere, & McDermott, 2018) and language processing (Caucheteux, Gramfort, & King, 2023). In the math domain, a Transformer-based ANN model has been shown to predict brain activity of adult participants when they solve a variety of arithmetic problems (Nakai & Nishimoto, 2023). Much like with complex visual shapes, deeper ANN layers showed a better fit to the activity patterns of mathematical operators, suggesting that the latent features underlying mathematical operations reflect higher-order information integrated through cumulative transformations of stimulus input. However, this study only included adult participants, who represent the end point of arithmetic learning. It remains unknown whether an ANN model may also predict arithmetic-related activity in children who are still in the process of learning arithmetic, and may therefore rely to a greater extent on counting strategies. Clearly, associationist models of arithmetic learning (Ashcraft, 1992; Chen & Campbell, 2018; Logan, 1988) would hypothesize that the performance of an ANN model in predicting arithmetic-related activity would improve with age and degree of exposure with simple problems, as associations between operands and answers would strengthen (**Figure 1A**).

**Figure 1.**
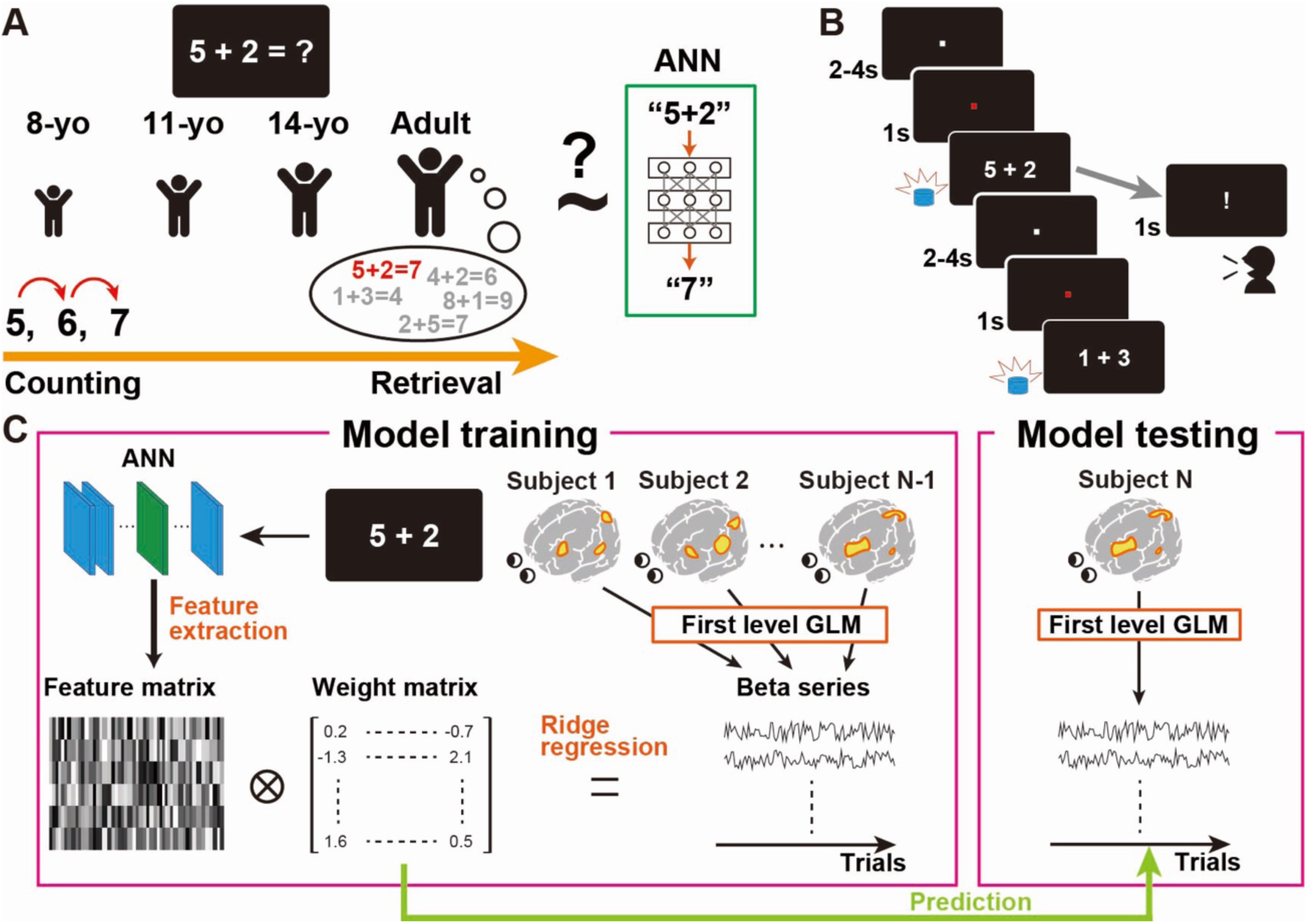
Hypothesis, experimental design and analyses. (**A**) Associative models of arithmetic learning assume that the decrease in overt counting strategies observed across development is explained by an increase in associations between operands of problems and answers, which would then be retrieved (see main text). Therefore, these models predict that solving arithmetic problems in older children and adults should be comparable to the way artificial neural networks (ANNs) provide the answer of an arithmetic problem. (**B**) In the scanner, participants were presented with single-digit addition problems and pressed a response button when they came up with an answer in their head (timing was self-paced). In some trials, participants said the answer aloud after being presented with an exclamation point. (**C**) Trial-wise brain signals were estimated for each subject using a general linear model (GLM). Vertex-wise encoding models were trained using feature matrices extracted from addition problems via ANNs and trial-wise brain signal estimates concatenated across participants. Prediction performance was evaluated with the data of the left-out subject.

To test this hypothesis, we measured fMRI activity of 104 participants in four age groups (8-year-olds [yo], 11-yo, 14-yo, and adults) while they solved single-digit addition problems (**Figure 1B**). We then compared arithmetic-related activity with ANN features using vertex-wise encoding models (Naselaris, Kay, Nishimoto, & Gallant, 2011). Encoding models, which are widely used in studies comparing brain activity and various ANN models, predict brain activity by a weighted linear combination of features extracted from experimental stimuli (Goldstein et al., 2022; Güçlü & van Gerven, 2015; Horikawa & Kamitani, 2017; Khosla, Ngo, Jamison, Kuceyeski, & Sabuncu, 2021; Nakai & Nishimoto, 2023). To mitigate effects of differences in processing times between different age groups, we first obtained trial-by-trial beta estimates of brain activity by means of a general linear model for each participant. We then built a series of vertex-wise encoding models predicting brain activity (trial beta estimates) from latent features extracted from a pre-trained large language ANN model (MathBERT) (Peng et al., 2021) (**Figure 1C**). The encoding model was trained for each age group and tested using leave-one-subject-out cross validation.

This computational approach enabled us to test two main hypotheses. First, if operands become increasingly associated with answers of problems over the course of learning and exposure to arithmetic problems, brain activity associated with simple addition problem-solving should become increasingly comparable to the way an ANN model provides the result of an arithmetic problem. Therefore, the ANN-based encoding model should show higher prediction performance for older participants (i.e., 14-yo and adults) than younger participants (i.e., 11-yo and 8-yo). Second, associationist accounts often predict that the strength of associations between operands and answers depends on the frequency of exposure (Ashcraft, 1992; Zbrodoff, 1995), which is believed to be higher for problems with a small sum than problems with a large sum (the latter being found less frequently in school manuals; (Hamann & Ashcraft, 1986)). Therefore, the ANN-based encoding model should show better prediction performance for relatively small addition problems (i.e., problems with operands ≤ 4) than problems that are larger in comparison (i.e., problems with operands ≥ 5).

## Results

### ANN better predicts arithmetic-related activity in older than younger participants in the left frontal cortex

To test whether ANN features may better predict brain activity during addition problem solving in older than younger participants, we independently constructed an encoding model (hereafter referred to as “ANN model”) and quantified model performance for each of the four age groups. Based on the observation in our previous study (Nakai & Nishimoto, 2023), we expected that deeper ANN layers would capture higher-order information related to mathematical operations than shallower layers and thus extracted ANN features from the last layer of the MathBERT. In adults, the ANN model significantly predicted activity in several brain regions, including the left precentral sulcus (L. PreCS), left precentral gyrus (L. PreCG), left insula, right middle frontal gyrus (R. MFG), and right postcentral gyrus (R. PostCG) (Wilcoxon sign-rank test: peak level, uncorrected *P* < 0.001; cluster level, corrected *P* < 0.05) (**Figure 2A**). In 14-yo, prediction accuracy was significant in the L. PreCS, as well as in the left occipital pole (L. OP), right anterior cingulate cortex (R. ACC), right superior temporal sulcus (R. STS), and left middle occipital gyrus (R. MOG) (**Figure 2B**). In 11-yo, the ANN model predicted activity in the L. MOG and right intraparietal sulcus (R. IPS) (**Figure 2C**). Finally, in 8-yo, no brain regions showed significant prediction accuracy (**Figure 2D**). Critically, the number of significant vertices increased with age (adults, 121 vertices; 14-yo, 92 vertices; 11-yo, 43 vertices; 8-yo, 0 vertices), even without cluster thresholding (adults, 391 vertices; 14-yo, 335 vertices; 11-yo, 219 vertices; 8-yo, 236 vertices) Therefore, the ANN model appeared to predict activity in larger brain regions in older participants.

**Figure 2.**
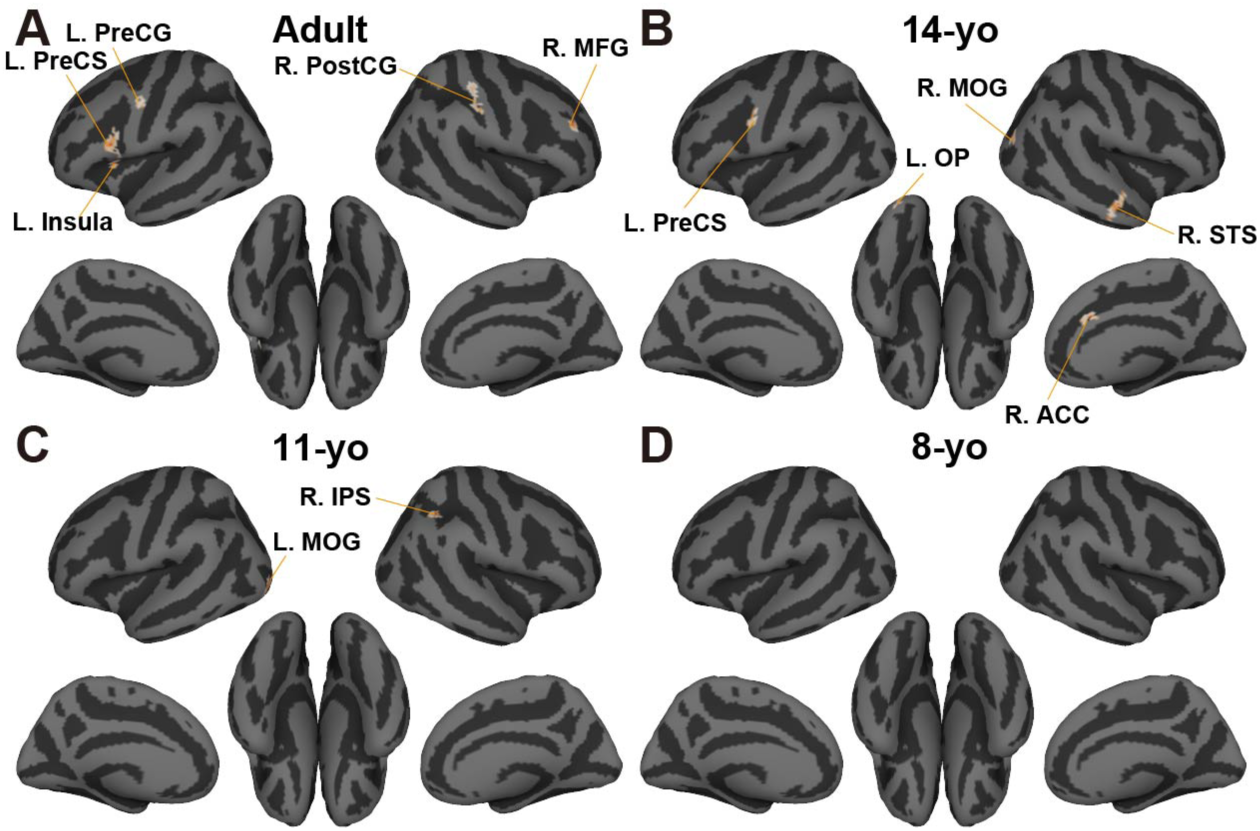
Prediction performance of ANN encoding model for each age group. Prediction accuracy of ANN encoding model using the 12^th^ layer mapped onto the cortical surface for (**A**) adults, (**B**) 14-yo, (**C**) 11-yo, and (**D**) 8-yo. Only significantly predicted vertices are shown (peak-level, uncorrected *P* < 0.001; cluster-level, false-discovery rate [FDR] corrected *P* < 0.05).

We then formally tested whether the prediction performance of the ANN model increased with age (and therefore frequency exposure to arithmetic problems). We first examined whether some brain regions displayed different prediction accuracy between the four age groups using a Kruskal-Wallis test (peak level, uncorrected *P* < 0.001; cluster level, corrected *P* < 0.05). To ensure that any between-group effect would only be observed in brain regions that also showed significant prediction accuracy in at least one of the groups, we inclusively masked that contrast by brain regions showing significant prediction accuracy in at least one of the four age groups (i.e., the union of **Figure 2A-D**). We found a significant main effect of age in the anterior and posterior L. PreCS (**Figure 3A**), indicating that prediction performance of the ANN model differed between groups in these two prefrontal clusters. Using post-hoc paired tests between groups, we then compared prediction accuracy by the ANN model for all combinations of age groups using functional regions-of-interest (ROIs) in the anterior and posterior PreCS. The two functional L. PreCS subclusters showed distinct patterns. In the anterior L. PreCS subcluster, which was located on the boundary of the opercular part of left inferior frontal gyrus (L. IFG), prediction accuracy was significantly larger in adults than in the other three groups (Wilcoxon rank-sum test: Adult vs. 14-yo, *P* < 0.001; *d* = 1.85; Adult vs. 11-yo, *P* < 0.001, *d* = 1.60; Adult vs. 8-yo, *P* = 0.001, *d* = 0.90; **Figure 3C**). In the posterior L. PreCS subcluster, both adults and 14-yo showed larger prediction accuracy than 11-yo and 8-yo (Adult vs. 11-yo, *P* < 0.001, *d* = 1.48; Adult vs. 8-yo, *P* = 0.003, *d* = 0.82; 14-yo vs. 11-yo, *P* < 0.001, *d* = 1.90; 14-yo vs. 8-yo, *P* < 0.001, *d* = 1.15; **Figure 3D**). Finally, a L. PreCS ROI defined anatomically showed a significant main effect of age (Kruskal-Wallis test, *P* = 0.002, *η^2^* = 0.161) and larger prediction accuracy in Adult and 14-yo than 11-yo (Adult vs. 11-yo, *P* < 0.001, *d* =1.03; 14-yo vs. 11-yo, *P* < 0.003, *d* =0.98; **Figure 3E**; also see **Figure S1** for other anatomical ROIs). These results suggest that the ANN model better predicts arithmetic-related activity in older than younger participants in the L. PreCS.

**Figure 3.**
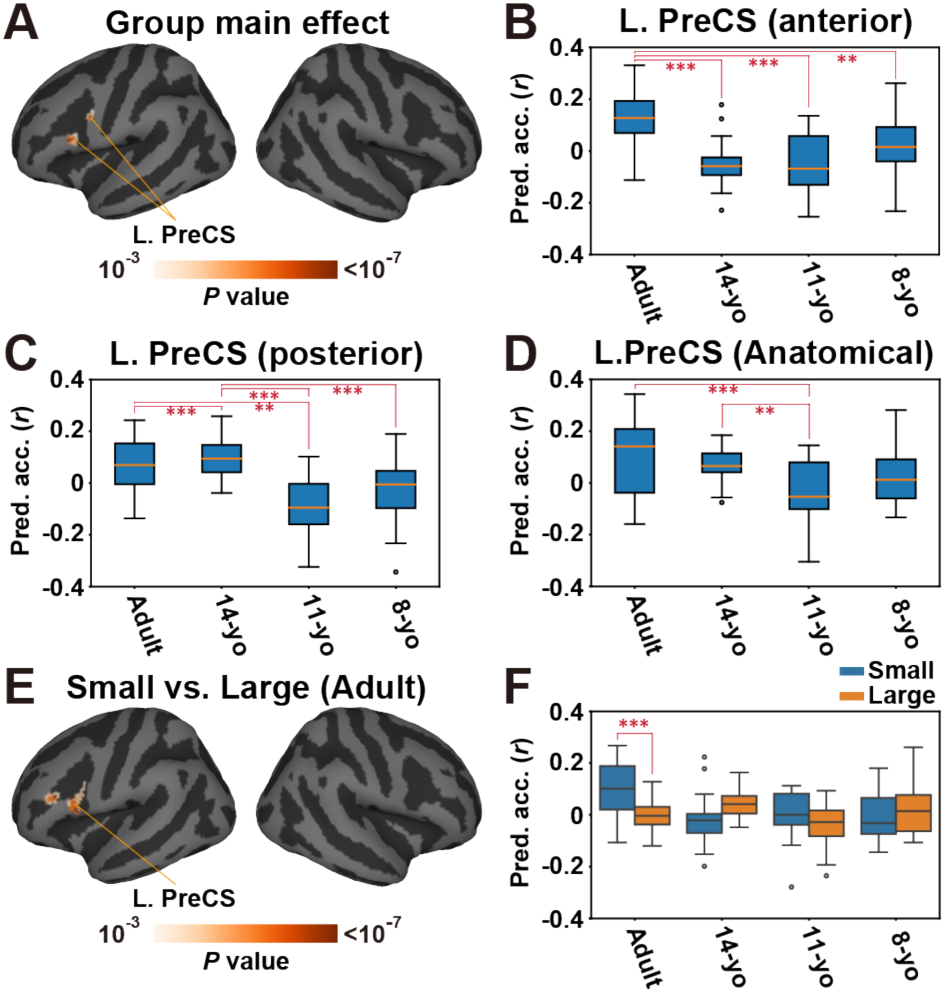
ANN prediction performance increases with age and decreases with problem size in the left PreCS. (**A**) A whole cortical map of the main effect of prediction accuracy across four age groups (Kruskal-Wallis test), within the inclusion mask of the union of **Figure 2A-D**. Statistical significance is evaluated with a peak-level *P* < 0.001 and cluster-level *P* < 0.05 (FDR corrected). (**B-D**) Box plots show the prediction accuracy of four age groups using the ANN model, averaged in the (**B**) anterior and (**C**) posterior parts of the L. PreCS functional ROI which are extracted from **Figure 3A**, and (**D**) anatomical L. PreCS ROI. (**E**) A whole cortical map of the direct comparison of prediction accuracy between small and large problems (Wilcoxon sign-rank test) for adults (see **Figure S3** for other age groups). (**F**) A box plot shows prediction accuracy of four age groups, separately calculated for small ≤ 4) and large (≥ 5) problems, averaged in the anatomical L. PreCS ROI. Error bar, SD.

One concern in evaluating between-age differences is the potential impact of differences in signal-to-noise ratio (SNR). Indeed, the higher prediction accuracy in the older group might be attributed to their higher SNR. To exclude this possibility, we calculated noise ceiling (i.e., maximum prediction accuracy) and signal-to-noise ratio values for each subject and compared them between different age groups (Schoppe, Harper, Willmore, King, & Schnupp, 2016) (**Figure S2**). In either index, we did not find any significant group differences for the whole-cortex comparison or the anatomical ROI analysis in the L. PreCS (noise ceiling, *P* = 0.849, *η^2^* = 0.009; SNR, *P* = 0.845, *η^2^* = 0.007), suggesting that the higher prediction accuracy of older groups was not caused by a reduced noise level.

### ANN better predicts arithmetic-related activity for smaller than larger problems in the left frontal cortex of adult participants

Next, we tested whether prediction performance was larger for small problems (where the operands are ≤ 4) than for large problems (where the operands are ≥ 5). Across the whole cortex, the contrast revealed significantly better prediction accuracy for small than large problems in the L. PreCS and left inferior frontal sulcus in adult participants, but not in any other groups (**Figures 3E, Figure S3**). An anatomical ROI analysis in the L. PreCS further replicated this better prediction accuracy for small than large problems in adults (Wilcoxon signed-rank test, *P* < 0.001, *d* = 1.26; **Figures 3F**). The difference between small and large problems was greater in adults than in other groups (Adult vs. 14-yo, *P* < 0.001, *d* = 1.52; Adult vs. 11-yo, *P* = 0.017, *d* = 0.63; Adult vs. 8-yo, *P* < 0.001, *d* = 1.07). These results suggest that the ANN model could better predict activity associated with small problems compared to activity associated with large problems, but only for adults who were the most exposed to arithmetic problems.

### ANN performance correlates with that of a model in which representations of individual addition problems are discrete in older participants

The ANN model consists of a large number of parameters, and its internal representation is often viewed as a “black box” (Ras, Xie, Van Gerven, & Doran, 2022; van der Velden, Kuijf, Gilhuijs, & Viergever, 2022). To shed some light on the reasons for an age-dependent increase in prediction accuracy of the ANN model, we performed several additional analyses. The first of these analyses assessed the correlation of prediction patterns across participants between ANN and different features. Specifically, we constructed additional encoding models using different features and compared their prediction patterns against those of ANNs. We reasoned that, if the ANN model exhibits similar prediction patterns as encoding models of other features, the ANN should represent information in a way that is similar to these features.

We defined three models. The ‘discrete’ model included features consisting of 45 one-hot vectors (i.e., 45 dimensions) representing all 45 addition problems as discrete instances. The ‘problem-size’ model included a one-dimensional feature corresponding to the sum of addition problems. Finally, the ‘difficulty’ model included a one-dimensional feature corresponding to reaction times of addition problems (averaged across all runs for each participant).

We found similar prediction patterns between the ANN model and the discrete model. The discrete model significantly predicted brain activity in the left PreCS for adults and 14-yo (**Figure S5A-B**), but only the left occipital cortex was significant for 11-yo (**Figure S5C**). No significant regions were found for 8-yo (**Figure S5D**). Participants showing higher prediction performance with the ANN model also exhibited higher prediction performance with the discrete model across the cortex (Spearman’s correlation coefficient, 0.621 ± 0.070; **Figure 4A**) and in the L. PreCS (*ρ* = 0.60; **Figure 4D**).

**Figure 4.**
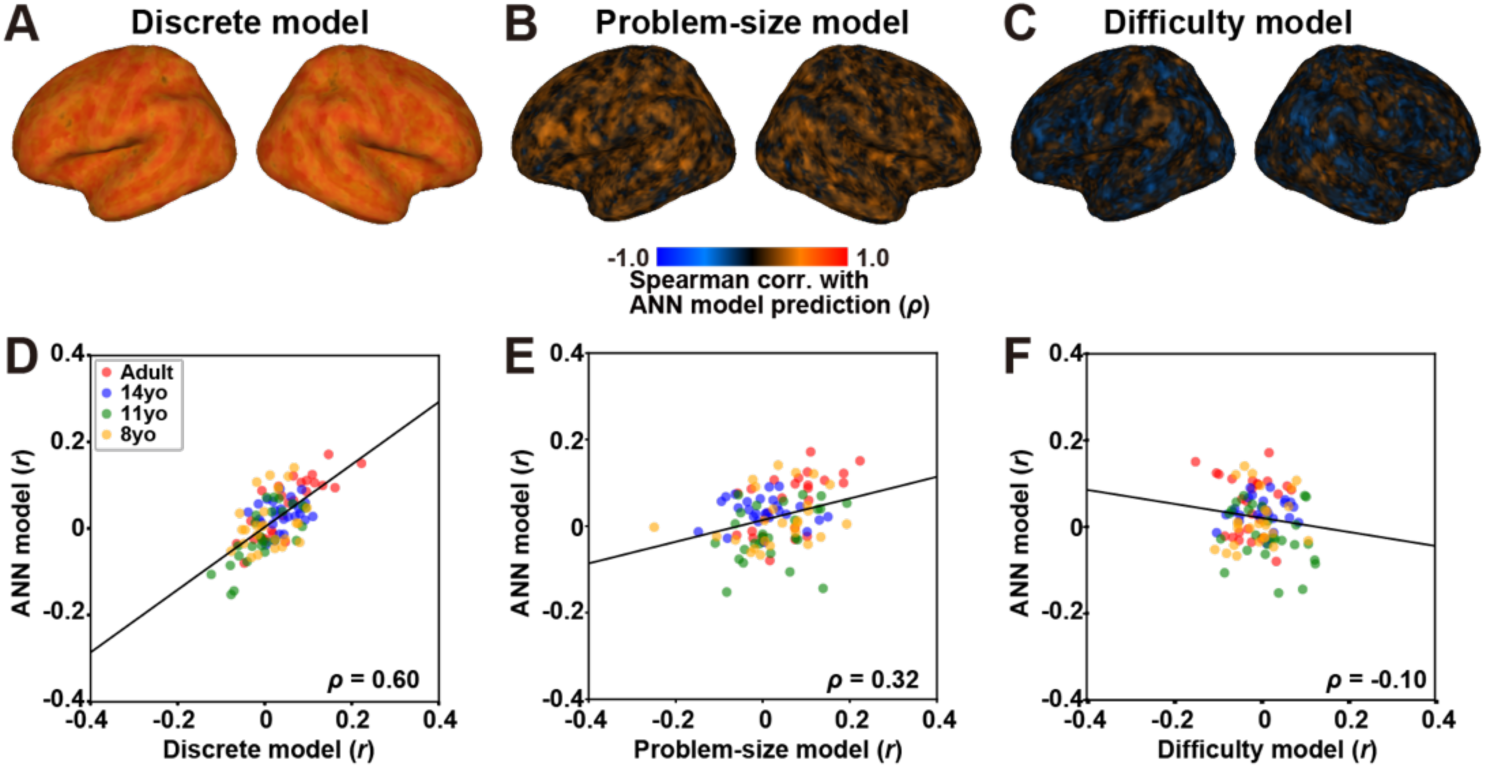
The discrete model exhibited prediction patterns that were similar to the ANN model. (**A-C**) Whole cortical maps show Spearman’s correlation coefficients of prediction accuracy (**A**) between the discrete and ANN models, (**B**) between the problem-size and ANN models, and (**C**) between the difficulty and ANN models. (**D-F**) Scatter plots show correlations of prediction accuracies (**D**) between the discrete and ANN models, (**E**) between the problem-size and ANN models, and (**F**) between the difficulty and ANN models, averaged in the left PreCS (anatomical ROI). The regression line and Spearman’s correlation coefficient (*ρ*, indicated at the bottom of each plot), are calculated based on the concatenated data across adults (red dots), 14yo (blue dots), 11yo (green dots), and 8yo (yellow dots).

In contrast, the problem-size and difficulty models displayed distinct patterns (**Figure S6, S7**). The mean prediction accuracies in the L. PreCS displayed a smaller correlation between the ANN and problem-size models (across the cortex, 0.148 ± 0.109; in L. PreCS, *ρ* = 0.32; **Figure 4B, E**) and between the ANN and difficulty models (across the cortex, 0.007 ± 0.111; in L. PreCS, *ρ* = -0.10; **Figure 4C, F**), indicating that problem-size information and cognitive load cannot explain prediction patterns of the ANN model. Overall, these results suggest that, in the ANN model, addition problems are more likely represented as discrete categories.

### Addition problems are more discretely represented in older participants

A second analysis aiming to shed some light on the reasons for an age-dependent increase in prediction accuracy of the ANN model involved a visualization of 45 addition problems using ANN model weights. Specifically, we visualized the relation among 45 addition problems using principal component analysis (PCA). First, we applied PCA to the discrete model weights (concatenated across four age groups) in the L. PreCS, and mapped all addition problems on the two-dimensional space using the first and second PC loadings.

We found that addition problems are most widely scattered in this space for adults (**Figure 5A**). Such representational dispersion decreased in younger participants: 14-yo, 11-yo, and 8-yo (**Figure 5B, 5C, 5D**, respectively). A quantitative assessment of weight variance showed a similar trend. The weight variance was significantly larger in older groups than in younger groups (Kruskal-Wallis test: *P* < 0.001, *η^2^* = 0.639; Wilcoxon rank-sum test: Adult vs. 14-yo, *P* < 0.001, *d* = 1.81; Adult vs. 11-yo, *P* < 0.001, *d* = 2.14; Adult vs. 8-yo, *P* < 0.001, *d* = 2.52; 14-yo vs. 11-yo, *P* < 0.001, *d* = 1.11; 14-yo vs. 8-yo, *P* < 0.001, *d* = 2.94; 11-yo vs. 8-yo, *P* < 0.001, *d* = 1.66; **Figure 5E**). Furthermore, the small problems (i.e., problems with operands ≤ 4) were more widely scattered compared to the large problems (Adult, *P* < 0.001, *d* = 1.45; 14-yo, *P* < 0.001, *d* = 1.16; 11-yo, *P* < 0.001, *d* = 0.86; 8-yo, *P* < 0.001, *d* = 0.70; **Figure 5F**), which is consistent with the prediction performance result (shown in **Figures 3F**). These results indicate that, for older groups, addition problems are represented as discrete categories in the brain. Such discrete representations could have contributed to prediction patterns in the discrete model (and hence in the ANN model too).

**Figure 5.**
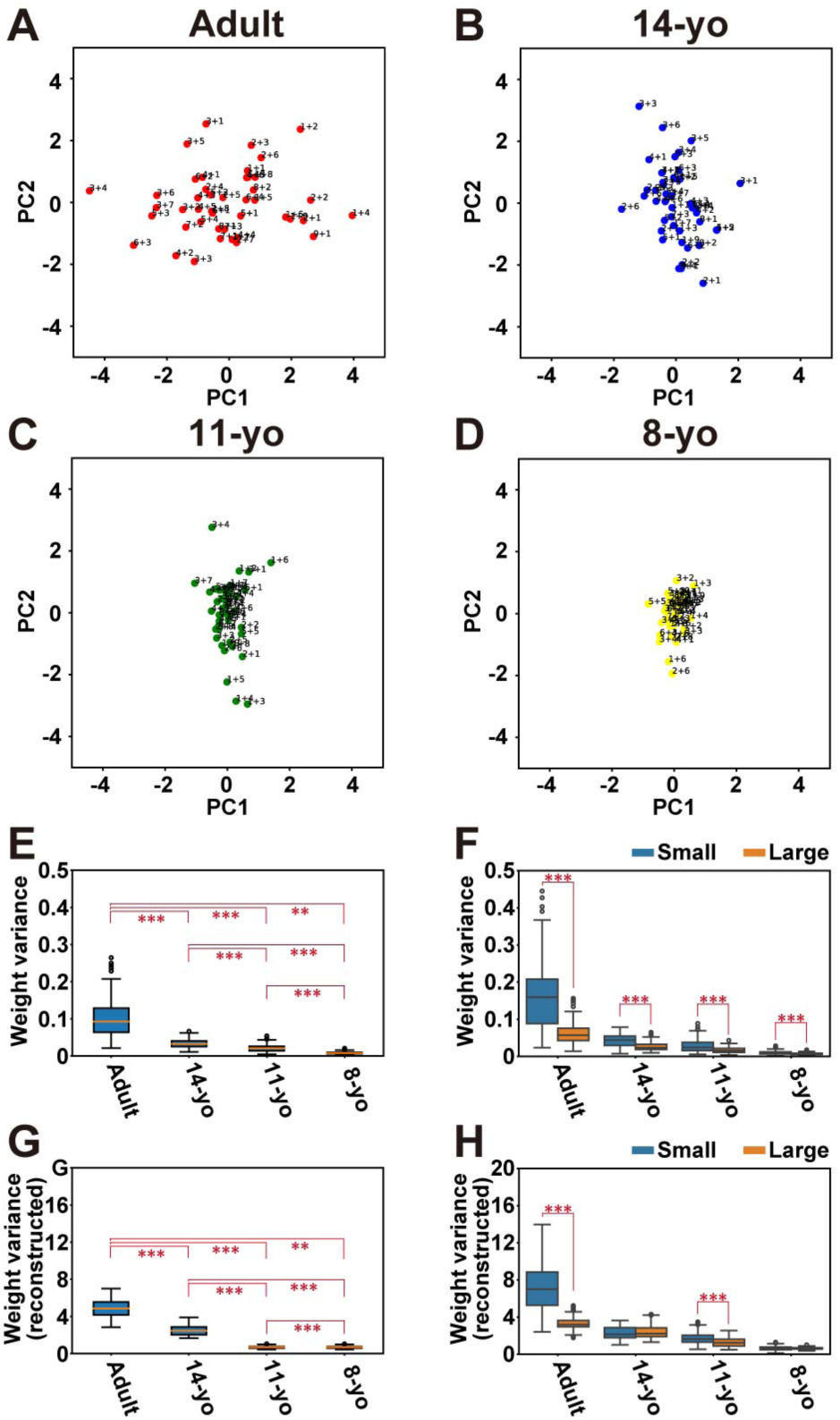
Representations of addition problems become discrete in older participants. (**A-D**) Principal component analysis based on the weight matrix of the discrete model. 45 addition problems are mapped onto the two-dimensional space of the first and second principal components (PC1 and PC2, respectively) based on their loadings, separately visualized for (**A**) adults, (**B**) 14yo, (**C**) 11yo, and (**D**) 8yo participants. (**E**) The variance of model weights across 45 addition problems is plotted using box plots. (**F**) The variance of model weights is separately calculated for small (problems with operands ≤ 4) and large (problems with operands ≥ 5) problems and plotted using box plots. (**G**) Discrete model weights are reconstructed using the ANN model, and the variance of the reconstructed weights across 45 addition problems is plotted using box plots. (**H**) The variance of reconstructed weights is separately calculated for small (problems with operands ≤ 4) and large (problems with operands ≥ 5) problems and plotted using box plots.

Next, we tested whether such discrete representations of addition problems spontaneously emerge using the ANN model without assuming discrete representations in the model. To this end, we reconstructed discrete model weights by multiplying the ANN features for each addition problem to the ANN model weights, resulting in highly similar reconstructed weights across the cortex (**Figure S8A-B**). We then performed PCA using the reconstructed weight values. We again found that addition problems are most widely scattered in the two-dimensional space for the adults than younger age groups (**Figure S8C-F**). The weight variance was significantly larger in older groups than in younger groups (**Figure 5G**), and the small problems were more widely scattered compared to the large problems (**Figure 5H**). These results indicate that addition problems are represented discretely within the ANN model, and such representations become more apparent with older age groups having more exposure to arithmetic problems.

### ANN predicts arithmetic-related activity in a layer-dependent manner

A third analysis aiming to shed light on the reasons for an age-dependent increase in prediction accuracy of the ANN model involved a comparison of prediction performance across different intermediate ANN layers. Indeed, our previous study had revealed a stronger association between arithmetic-related activity and ANN features in deeper layers (Nakai & Nishimoto, 2023), which is in line with a large number of studies investigating association between ANNs and human brain (Caucheteux et al., 2023; Cichy et al., 2016; Güçlü & van Gerven, 2015; Horikawa & Kamitani, 2017; Kell et al., 2018). If arithmetic-related representations in older participants are more similar to the ANN model than in younger participants, such hierarchical correspondence would be more evident in older than younger participants.

To test this possibility, we extracted latent features from the shallowest layer (i.e., first hidden layer) to the deepest layer (twelfth hidden layer) and examined whether such layer-dependent prediction patterns depend on age groups. We observed a topographic expansion of significant vertices in the L. PreCS for adult participants (**Figure 6A-B**). That is, deeper ANN layers predicted larger brain regions for this age group. Such topographic expansion in the frontal cortex was absent in younger groups (**Figure 6C-I**). In contrast, there was a topographic reduction of significant vertices in the left IOG in adults and 14-yo (**Figure 6A-E**). The same pattern was observed in 11-yo and 8-yo, but in a different sub-portion of the occipital cortex (i.e., L. LG) (**Figure 6F-I**). In sum, prediction performance by the ANN model increased with intermediate layers in the left frontal region in adults and 14-yo and in the right parietal region in adults, 11-yo, and 8-yo. In contrasts, it decreased in the occipital region in all age groups.

**Figure 6.**
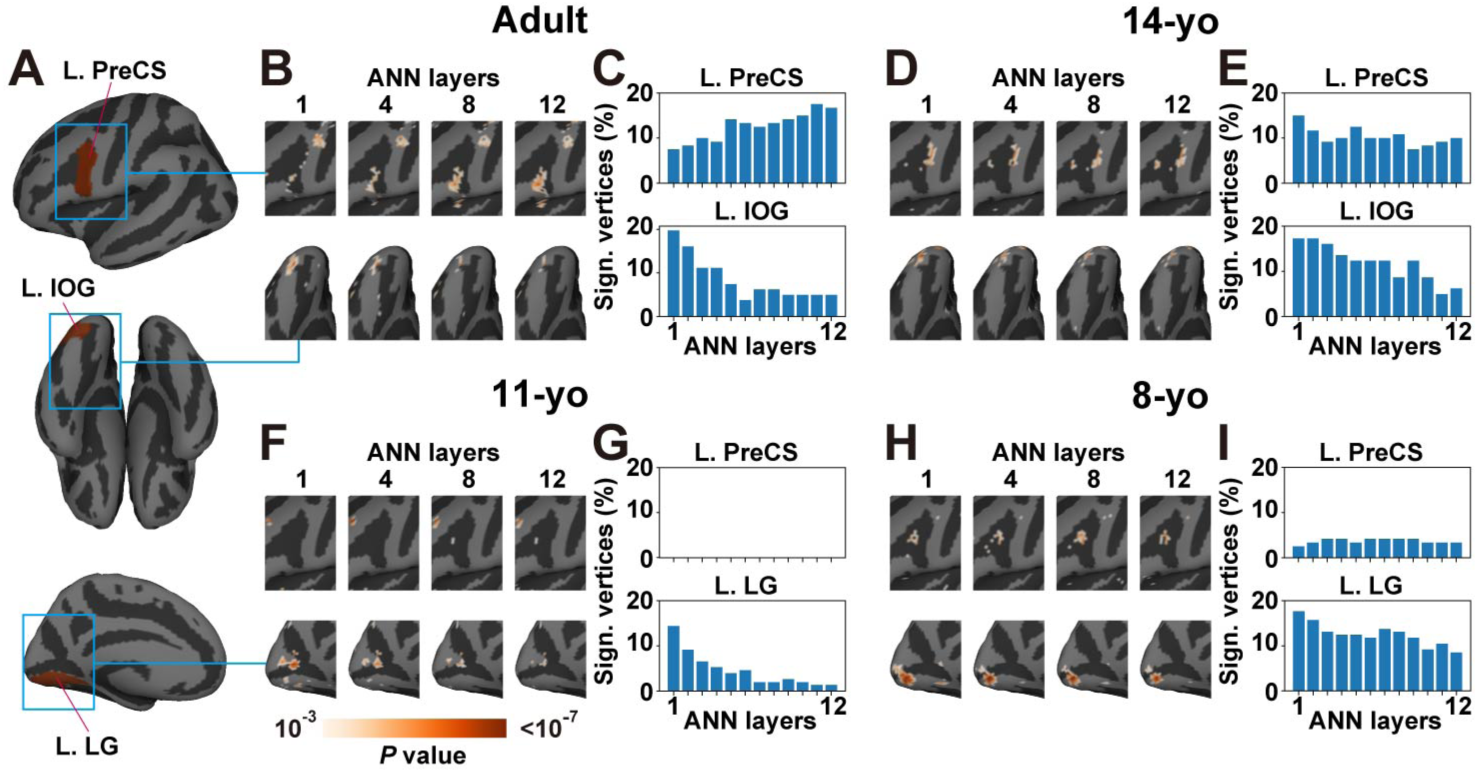
Topographic expansion and reduction of predictable brain regions with ANN layers across age groups. (**A**) The target anatomical regions-of-interest (in red) and the surrounding areas (in blue squares) of the left precentral sulcus (L. PreCS), left inferior occipital gyrus (L. IOG), and left lingual gyrus (L. LG) are shown on the cortical surface. (**B, D, F, H**) Prediction patterns are shown for the 1^st^, 4^th^, 8^th^, and 12^th^ ANN layers. Only significantly predicted vertices are shown (peak-level, uncorrected *P* < 0.001; cluster-level, corrected *P* < 0.05). Areas around the L. PreCS and L. IOG are shown for (**B**) adults and (**D**) 14-yo, while those around the L. PreCS and L. LG are shown for (**F**) 11-yo and (**H**) 8-yo. (**C, E, G, I**) Bar graphs of the percent of significant vertices are plotted for all of the twelve ANN layers. Bar graphs of the L. PreCS and L. IOG are shown for (**C**) adults and (**E**) 14-yo, while those of the L. PreCS and L. LG are shown for (**G**) 11-yo and (**I**) 8-yo.

These observations suggest that between-age effect in the ANN model performance may depend on the target layers. Indeed, we found a significant main effect of age of prediction accuracy in the L. PreCS (Kruskal-Wallis test: peak level, uncorrected *P* < 0.001; cluster level, corrected *P* < 0.05) when using features extracted from the 8^th^ layer (**Figure S9**), but no significant effect was found when using features extracted from the 4^th^ and 1^st^ layers. This suggests that age-related representational changes in arithmetic emerge primarily in deeper layers of the ANN model.

## Discussion

Most current models of arithmetic processing assume that arithmetic learning involves an increase in memory-based retrieval of associations between operands of problems and their answers (Ashcraft, 1992). Because this process bears similarity with the way information is retrieved in ANNs (i.e., through retrieval of statistical associations between elements), we asked in the current study whether arithmetic-related brain activity would increasingly resemble brain activity predicted by an ANN-based encoding model with age. Measuring fMRI activity associated with single-digit mental addition in participants from age 8 to adulthood, we found that it was the case in a region of the L. PreCS, particularly for the smallest problems. Further analyses showed that ANN predictions best aligned with a model in which representations of arithmetic problems were discrete, compared to models in which problems were categorized as varying with problem-size or difficulty. PCA visualization revealed that addition problems were more discretely distributed in older than in younger participants, even when using discrete model weights reconstructed via ANN features. Finally, layer-dependent increase in prediction accuracy was primarily observed in the L. PreCS in adults. To some extent, these findings suggest that neural activity associated with mental arithmetic can be increasingly predicted by an ANN with learning and development. However, this was largely constrained to a specific area of the left frontal cortex.

The developmental increase in similarity between arithmetic-related activity and predictions from an ANN in the L. PreCS is consistent with the idea that, with increasing age, there is some degree of greater similarity between the way arithmetic problems are solved by human participants and ANNs. This is in keeping with associative models of arithmetic learning. Indeed, these models assume that operands and answers of problems are progressively associated with one another through repeated exposure (Ashcraft, 1992; Chen & Campbell, 2018), which reminds of statistical learning in ANNs. Interestingly, because smaller problems tend to be more frequently encountered than larger problems (Hamann & Ashcraft, 1986), associative models typically predict that associations between operands and answers decrease with problem size (Chen & Campbell, 2018). In line with this idea, we found that activity associated with the smallest problems (i.e., problems with operands ≤ 4) show greater similarity with activity predicted by the ANN-based model than large problems (i.e., problems with operands > 4) in older participants.

On the one hand, the similarity between arithmetic-related activity and predictions from an ANN provides some support for the idea that solving single-digit addition problems in adults and older children may rely on retrieving associations between operands and answers from memory, much like an ANN may have learned to associate co-occurring elements in its training set. On the other hand, our ANN model was only able to increasingly predict arithmetic-related activity with age in a cluster of the left frontal cortex, located in the L. PreCS. In other words, we did not find age-related increase in ANN prediction accuracy in most of the brain, including in regions that are classically associated with mental arithmetic such as the bilateral IPS, left AG, and hippocampus (Istomina & Arsalidou, 2024; Sokolowski, Hawes, & Ansari, 2023). A lack of similarity between ANN predictions and IPS activity may be explained by the fact that IPS-related activity in mental arithmetic is often interpreted as reflecting explicit manipulation of quantities rather than retrieval of associations (Zamarian, Ischebeck, & Delazer, 2009). However, the lack of similarity with the left AG and the hippocampus is more surprising, given the hypothesized role of these regions in memory-based retrieval of associations (Grabner et al., 2009; Klein, Willmes, Bieck, Bloechle, & Moeller, 2019; Qin et al., 2014; Sokolowski et al., 2023). In fact, we even found a decrease in prediction accuracy with age using the ANN model in the right hippocampus (**Figure S1**).

We can see two possibilities to explain the restricted similarity between the ANN model and arithmetic-related activity in the L. PreCS. First, it is possible that this region may play a more critical role in memory retrieval than regions such as the hippocampus (Sokolowski et al., 2023), which may support encoding of associations between operands and answers in the early stages of arithmetic learning but not retrieval of these associations later on (Qin et al., 2014). Indeed, in contrast to the hippocampus, activations in the left frontal cortex (**Figures 2, 3**) have frequently been observed in arithmetic task fMRI. Coordinate-based meta-analyses indicate that activation of the L. PreCS/PreCG and neighboring L. IFG is frequently observed in tasks involving addition problem solving (Arsalidou & Taylor, 2011; Istomina & Arsalidou, 2024). Activation in the L. IFG in arithmetic tasks also overlapped with sentence comprehension (Nakai & Okanoya, 2018, 2020) and semantic memory retrieval tasks (Heidekum, Grabner, De Smedt, De Visscher, & Vogel, 2019), which may suggest that this region contributes to the retrieval of arithmetic facts from memory. Overall, these observations are in line with developmental neuroimaging studies reporting brain-behavior correlations between arithmetic skills and structural properties of left frontal areas (Rotzer et al., 2008; Viesel-Nordmeyer & Prado, 2023), as well as anatomical connectivity between the L. PreCS/IFG and IPS (Tsang, Dougherty, Deutsch, Wandell, & Ben-Shachar, 2009). Note that our study identifies a cluster located in the L. PreCS rather than the L. IFG per se. However, because volume-based analysis may ignore information on cortical folding patterns and is less sensitive to local spatial patterns (Oosterhof, Wiestler, Downing, & Diedrichsen, 2011; Tucholka, Fritsch, Poline, & Thirion, 2012), we adopt here a surface-based analysis (Dale, Fischl, & Sereno, 1999; Fischl, Sereno, & Dale, 1999). Therefore, clusters in the dorsal and posterior parts of the IFG identified in previous volume-based analyses may be relatively close to the L. PreCS identified here.

There is a second possibility to explain the relatively restricted similarity between arithmetic-related activity and predictions from the ANN: Retrieval of associations from memory may not be the only path to simple arithmetic problem solving in adults and older children. Indeed, a series of recent studies (Barrouillet & Thevenot, 2013; Díaz-Barriga Yáñez et al., 2023; Fayol & Thevenot, 2012; Poletti, Díaz-Barriga Yáñez, Prado, & Thevenot, 2023; Poletti, Perez, Houillon, Prado, & Thevenot, 2021; Thevenot et al., 2016; Uittenhove, Thevenot, & Barrouillet, 2016) suggest that counting procedures that may be used by young children when learning to add numbers may be accelerated up to the point that they become automatic and unconscious (Uittenhove et al., 2016). This may explain a range of behavioral and neuroimaging findings, including the presence of a problem size effect with small addition problems (Uittenhove et al., 2016), the difference between single-digit addition and single-digit multiplication (Fayol & Thevenot, 2012; Mathieu, Epinat-Duclos, Léone, et al., 2018; Mathieu, Epinat-Duclos, Sigovan, et al., 2018), and the recent findings that patterns of brain activity associated with the problem size effect change quantitatively but not qualitatively over development (Díaz-Barriga Yáñez et al., 2023). Interestingly, it has recently been proposed that automatized counting may coexist (and compete) with retrieval of associations from memory (Prado, Thevenot, under review). Clearly automatized counting is vastly different from the type of statistical learning at the core of an ANN. Therefore, the coexistence of both automatized counting and retrieval of associations in the human mind during arithmetic problem-solving may explain the relatively limited accuracy of ANN predictions in the present study.

Interestingly, among the four models implemented in this study, the difficulty model was the only one to show greater similarity with brain activity patterns in younger than older participants (in the primary motor cortex) (**Figure S7**). We speculate that activity in this region may reflect some finger-related processing supporting counting. For example, previous studies reported activity of the primary motor and premotor cortices during counting tasks (Hinton, Harrington, Binder, Durgerian, & Rao, 2004; Piazza, Mechelli, Price, & Butterworth, 2006; Zago et al., 2010). Indeed, the difficulty model showed significant prediction in the dorsal part of the central sulcus (**Figure S7**), which is associated with finger movements rather than with the mouth and tongue (Gordon et al., 2023; Penfield & Boldrey, 1937). The high predictive accuracy of this model for the younger groups, as well as a tendency of negative correlation between the ANN and difficulty model (**Figure 4**), is consistent with the fact that younger participants may frequently adopt silent counting strategies (Ashcraft, 1982; Groen & Parkman, 1972; Qin et al., 2014; Wu et al., 2008).

To shed some light on the nature of the relation between the ANN model and brain activity, we performed several additional analyses comparing the ANN model to other models in which arithmetic problems were categorized in different ways. There was a positive correlation between prediction performance of the ANN model and prediction performance of a model in which addition problems were each represented separately, suggesting that the ANN model may represent (at least in part) each addition problem discretely (**Figure 4**). As we observed highly positive correlation across the entire cortex, this effect is likely due to the shared features underlying the ANN and that ‘discrete model,’ rather than to the specific brain activity pattern of the L. PreCS. PCA results further support these findings. Increasing weight variance (**Figure 5**), as well as their two-dimensional visualization, indicated that older participants have more discrete representations of each addition problem in the brain. Moreover, the ANN-based reconstruction analysis indicates that such discrete representations can be obtained using the ANN model, even using continuous latent features of addition problems. These results suggest that the ANN might have been able to predict arithmetic-related activity of older participants because their brains represented addition problems discretely, and the ANN model effectively captured such representations. Note, however, that the analyses above do not necessarily mean that the brain representations of addition problems can be entirely explained as discrete instances. Indeed, a model in which problems were categorized as a function of their size also showed a positive (though reduced) correlation with the ANN model (**Figure 4**), suggesting that the ANN model may contain some information about problem-size, which is well known to influence brain activity (Díaz-Barriga Yáñez et al., 2023).

The hierarchical prediction pattern in the left frontal and visual cortices (**Figure 6**) is in line with previous studies in visual neuroscience combining ANN and human data (Cichy et al., 2016; Güçlü & van Gerven, 2015; Horikawa & Kamitani, 2017), where deeper ANN layers showed increase in prediction performance in higher-order visual cortex. Such patterns have also been observed in the auditory (Kell et al., 2018; Tuckute, Feather, Boebinger, & McDermott, 2023), language (Caucheteux et al., 2023; Goldstein et al., 2022), and arithmetic processing in adults (Nakai & Nishimoto, 2023). In arithmetic processing, input information is thought to ascend the brain’s hierarchy from lower-level sensory areas to higher-level association cortex (Dehaene, Molko, Cohen, & Wilson, 2004; Vogel & De Smedt, 2021), in line with the observed decrease in prediction in areas of the visual cortex the increase in prediction in the left frontal cortex. This might indicate that deeper ANN layers hold higher-order information than simple visual features. More generally, sensory perception such as vision and audition and abstract cognitive functions such as language and mathematics may not follow exactly the same hierarchical pathways in the brain. However, the current study shows that there could be a common principle regarding the correspondence between ANNs and BNNs in these different domains.

We note three limitations of the current study. First, our encoding models did not directly predict brain activity but rather beta estimates after estimating first-level GLM for each participant. Although this approach deviates from conventional encoding models using naturalistic stimuli (e.g., movie viewing task) (Huth, Nishimoto, Vu, & Gallant, 2012; Nishimoto et al., 2011), it enables us to minimize the impact of performance (e.g., response times) variations from trial to trial and to compare model performance across age groups using the same training and testing sample size. Nevertheless, because this method involves fitting the data in two stages, it might downplay information about the dynamic processing of arithmetic in the first stage. Second, our task design contained only addition problems. Therefore, it remains unclear whether our results may apply to other arithmetic operations. Nonetheless, in a previous study, we have shown that an ANN encoding model may predict brain activity of addition, subtraction, multiplication, and division problems, as well as their combinations (e.g., “(3 + 2) × 4”) (Nakai & Nishimoto, 2023). Third, the ANN used in the current study (MathBERT) was trained with a very large dataset of mathematical problems, which contains not only simple arithmetic problems but also more complex algebra (Peng et al., 2021). This arguably departs from the way children acquire simple arithmetic skills. Future studies using more realistic models based on actual learning materials (i.e., math textbooks) may allow for more accurate models that could potentially predict brain representations of arithmetic in children.

## Conclusion

In summary, this study demonstrates that addition problem-solving in the brain increasingly aligns with ANN models with learning and development, particularly in the left precentral sulcus. Such an alignment is likely to arise from increasingly discrete organization of addition problems in the brain, especially for smaller problems. On the one hand, this generally supports associative models of arithmetic learning (e.g., Ashcraft, 1992). On the other hand, the limited correspondence between arithmetic-related activity and predictions from the ANN outside the precentral sulcus suggests that arithmetic processing in humans and ANNs is not entirely analogous. These findings provide insights into the neural and computational foundations of mathematical cognition, supporting the idea that the ability to learn simple arithmetic may only partly rely on associative memory mechanisms similar to those underlying ANNs, where repeated exposure strengthens the relations between operands and outcomes.

## Methods

### Participants

One hundred and forty-nine children, adolescents, and adults between the ages of 8 and 30 were recruited for the study (see also Díaz-Barriga Yáñez et al. (2023)). Participants were contacted through advertisements on social media. They were divided into four age groups: 8-to 9-year-olds (hereafter 8-yo, n = 41), 11-to 12-year-olds (hereafter 11-yo, n = 36), 14-to 15-year-olds (hereafter 14-yo, n = 30), and adults over the age of 18 (n = 42). Ten participants were excluded from the final analyses because they did not complete the fMRI session (8-yo, n = 4; 11-yo, n = 3; 14-yo, n = 3). Fourteen participants were also excluded after fMRI preprocessing, either because of excessive head motion in the scanner (see criteria below) or poor image quality (8-yo, n = 8; 11-yo, n = 2; 14-yo, n = 1; adults, n = 3). Finally, in order to match the number of participants in the four age groups, we randomly selected 26 participants (the minimum group size across the four groups) from each group. The final sample (n = 104) consisted of 26 participants in each of the four groups.

All participants were native French speakers. Participants below the age of 18 gave their assent to participate in the study, while their parents gave written informed consent. Adult participants gave written informed consent. The study was approved by a French ethics committee (Comité de Protection des Personnes Est 4, IDRCB 2019-A01918–49). Participants (or parents if participants were below 18) were paid 40 euros per session for their participation.

### Experimental procedure

The study consisted of two separate testing sessions: one behavioral session (which included psychometric and behavioral testing, as well as mock MRI scanning for children and adolescents) and one fMRI session.

In the first testing session, children and adolescents were familiarized with the fMRI environment in a mock scanner. They listened to a recording of the noises associated with all fMRI sequences. A motion tracker system (3D Guidance track STAR, Ascension Technology Corporation) was used to measure head movements and provide online feedback to the participants. Finally, participants practiced 25 trials of the silent arithmetic task in that mock scanner.

In the second testing session, participants performed a mental addition task in the MRI scanner. The task consisted of all 45 single-digit addition problems with a sum less than or equal to 10. Problems were presented in both commutative orders (e.g., 2 + 3 and 3 + 2). Each addition problem was presented five times over five consecutive runs to maximize signal-to-noise ratio in the fMRI scanner, for a total of 225 trials. In each trial, an addition problem (which remained on the screen until a response was detected) was preceded by a red square for a fixed duration of 1,000 ms and followed by a variable period of fixation (i.e., a white square) from 2,000 to 4,000 ms. Participants were instructed to not say the answer to each problem out loud but rather to press a button with their right hand as soon as they came up with the answer in their head. To ensure that participants performed mental calculations, some problems were immediately followed by an exclamation mark on the screen for 1,000 ms. In such cases, participants had to quickly say the answer out loud before the disappearance of the exclamation mark. Answers were recorded manually by the experimenter. There were four arithmetic problems of such kind for every run of 45 trials.

Trial order within each run and run order was randomized across the participants. Trials with vocal response were pseudo-randomized in the trial list, such that these trials were not located at the beginning of a run and were not consecutive. The task was programmed and presented with PsychoPy3 (v2020.2.5). Visual stimuli were projected on an in-scanner display by a projector in the room adjacent to the scanner. Head movement was minimized by cushions placed around the participant’s head.

### fMRI data acquisition

Images were collected using a Siemens Prisma 3 T MRI scanner with a 64-channel receiver head-neck coil (Siemens Healthcare, Erlangen, Germany) at the CERMEP Imagerie du vivant in Lyon, France. The blood oxygenation level-dependent (BOLD) signal was measured with a susceptibility-weighted single-shot echo planar imaging sequence. Imaging parameters were as follows: repetition time (TR) = 2,000 ms, echo time (TE) = 24 ms, flip angle = 80°, field of view (FOV) = 220 × 206 mm^2^, resolution = 1.72 × 1.72 mm^2^, slice thickness = 3 mm (0.48 mm gap), number of slices = 32. A high-resolution T1-weighted whole-brain anatomical volume was also collected for each participant. Parameters were as follows: TR = 2,400 ms, TE = 2.81 ms, flip angle = 8°, FOV = 224 × 256 mm^2^, resolution = 1.0 × 1.0 mm^2^, slice thickness = 1.0 mm, number of slices = 192.

### fMRI data preprocessing

The preprocessing of both functional and anatomical MRI data was carried out using fMRIPrep 21.0.2 (Esteban et al., 2019). The T1-weighted image underwent a correction for intensity non-uniformity using ANTs 2.3.3 N4BiasFieldCorrection (Avants et al., 2011) and was skull-stripped using a Nipype implementation of the antsBrainExtraction.sh workflow. Brain tissue segmentation was conducted using fast (FSL 6.0.5.1) (Y. Zhang, Brady, & Smith, 2001). Brain surfaces were reconstructed using recon-all (FreeSurfer 6.0.1) (Dale et al., 1999). Volume-based spatial normalization to the standard space (MNI152NLin6Asym) was achieved through nonlinear registration with antsRegistration (ANTs 2.3.3).

For the functional scans, a reference volume and its skull-stripped version were initially produced using the custom methodology of fMRIPrep. Prior to spatiotemporal filtering, head-motion parameters, including six rotation and translation parameters, were estimated in relation to the reference scan using mcflirt (FSL 6.0.5.1) (Jenkinson, Bannister, Brady, & Smith, 2002). All functional scans underwent slice-time correction to the middle slice using 3dTshift from AFNI (Cox & Hyde, 1997). The reference scan was subsequently co-registered with the T1w reference using bbregister (FreeSurfer) (Greve & Fischl, 2009). Three global signals were extracted within the cerebrospinal fluid (CSF), white matter (WM), and whole-brain masks. The blood oxygenation level-dependent time-series were resampled into the standard spaces (MNI152NLin6Asym). Runs in which translations greater than 2 mm or rotations greater than 2 radians were detected in more than 5% of volumes were excluded from the analysis (8-yo, n=3; 11-yo, n=2; 14-yo, n=1 of runs). Participants having less than three available runs were excluded (see Participants subsection).

The resulting functional data underwent high-pass filtering at 0.01 Hz and low-pass filtering at 0.15 Hz. Confounding factors, including the six head-motion parameters, global signals from the CSF, WM, and whole-brain masks, as well as the derivative of each factor, were eliminated using linear regression implemented in Scikit-learn. The functional data were then standardized to have a zero mean and unit variance for each of the 18,715 cortical vertices in the fsaverage5 space, excluding non-cortical vertices. Spatial smoothing was applied using a Gaussian filter with 8 mm full-width at half-maximum based on the geodesic distance on the cortical surface.

### Trial-based signal estimation

First-level individual analysis was performed based on the preprocessed fMRI data using the Nilearn package in Python (Abraham et al., 2014). Univariate activity (i.e., beta value) associated with the 45 mental addition problems was analyzed using a general linear model, where the fMRI signal for each problem was modeled as an epoch of the same length as the reaction time, starting with the presentation of the problem, convolved with a hemodynamic response function. In addition, we included the following regressors of non-interest: a button response regressor with a duration of 0.5 s starting with each button response, three drift regressors describing low-frequency signals, regressors representing experimental runs, and a constant regressor. Forty-five beta series representing brain activity associated with each arithmetic problem were obtained for each of the 104 participants (i.e., 26 participants for each of the four age groups).

### Feature extraction

Latent ANN features were extracted using a pre-trained large language model specifically tailored for understanding mathematical formulas, MathBERT (https://huggingface.co/tbs17/MathBERT) (Peng et al., 2021). Each addition problem was used as input, and intermediate outputs from the first to the twelfth layers (out of a total of twelve layers) were extracted as features. These resultant series of features were further reduced to 100 dimensions using PCA.

Latent features from ANN models were compared to encoding models that incorporated three features that represented different aspects of the problems: ‘discrete’ features consisting of 45 one-hot vectors (i.e., 45 dimensions) representing all 45 addition problems as distinct categories, a ‘problem-size’ feature corresponding to the sum of addition problems, and a ‘difficulty’ feature corresponding to reaction times of addition problems (averaged across all runs for each participant).

### Vertex-wise encoding model fitting

In the encoding model, each vertex’s cortical activity (beta series concatenated across participants, denoted as **R** of dimensions [T × V]) was modeled by multiplying the feature matrix **F** of dimensions [T × N] by the weight matrix **W** of dimensions [N × V] (where T = number of samples; N = number of features; V = number of vertices):

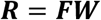

Model performance was evaluated using a nested leave-one-out cross-validation across participants. The weight matrix **W** was obtained using L2-regularized linear regression with the training data of 25 participants, which consisted of 1,125 samples. The optimal regularization parameter was determined using 10-fold cross-validation in the inner loop, where the regularization parameters varied from 1 to 10^6^.

The test data of the left-out participant consisted of 45 samples for each outer loop. The prediction accuracy was evaluated using Pearson’s correlation coefficient between the predicted and actual test samples. The peak-level statistical significance (one-sided) was calculated using a Wilcoxon rank-sum test (*p* < 0.001). A null distribution of cluster size was assessed with 1,000 permutation samples, and cluster-level significance was calculated based on this null distribution (*p* < 0.05), corrected for multiple comparisons using the false-discovery rate (FDR) (Benjamini & Hochberg, 1995). Pycortex was used to visualize data on cortical maps (Gao, Huth, Lescroart, & Gallant, 2015).

### Principal component analysis

Brain representations for the 45 addition problems were visualized using PCA. For this analysis and the following feature-brain similarity (FBS) analysis, we calculated a weight matrix using data from all participants (without leaving out one participant) into model training for each age group. To control the effect of varying scales of the weight matrices, we selected the same regularization parameter across all participant groups (i.e., the mode of the four age groups). We selected the vertices included in the anatomical L. PreCS ROI. We applied PCA to the [45*V] weight matrices (V = number of vertices, concatenated across four groups) of the encoding model. To show the structure of the representational space of addition problems, 45 problems were mapped onto the two-dimensional space using the loadings of the first and second PCs as the x-axis and y-axis, respectively.

### Weight reconstruction

To bridge the discrete model and ANN model weights, we reconstructed weight values of 45 addition problems of the discrete model using the ANN model weights. We first extracted latent ANN features of each addition problem (here denoted as *reference feature vectors*) and multiplied these features with the weight vectors obtained from the ANN encoding model. This analysis highlights how well the latent features can mediate between categorical concepts (addition problems in this case) and the brain. As a result, we obtained a 45-dimensional vector (i.e., reconstructed weights) in each vertex, which can be interpreted as a reconstruction of the discrete model weights via ANN latent features. We then performed a PCA using these reconstructed weights

### Noise ceiling and signal-to-noise ratio

To compare the signal-to-noise level of different age groups, we applied a *noise ceiling* calculation (Schoppe et al., 2016). Due to a need for repeated signals for this calculation, we divided the original fMRI data into two subsets: the first to second runs, and the third to fourth runs (the fifth run was not used in this analysis). A general linear model was fitted for each of these subsets, resulting in two sets of beta series for each subject.

The noise ceiling is based on the formula introduced by Sahani and Linden (2002). First, the variance of repetitive neural responses (i.e. total power; TP) is split into estimators of signal power (SP) and noise power (NP):

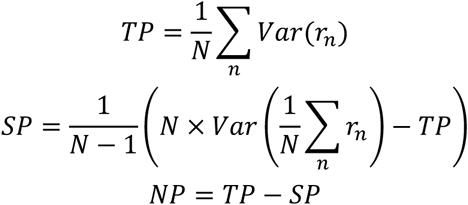

Where *N* is the total number of repetitions (= 2 in the current study), *r_n_* is a time series of neural responses at the *n*th repetition. Based on these estimators, the noise ceiling value *NC* (maximum prediction accuracy) can be calculated as follows:

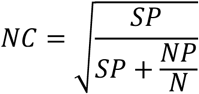

In the present study, we further calculated the signal-to-noise ratio (SNR) by dividing the signal power by the noise power:

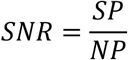

## Author contributions

**T.N.**: Conceptualization, Formal analysis, Visualization, Writing-Original draft preparation, Funding acquisition. **J.P.**: Supervision, Writing - Review & Editing, Funding acquisition.

## Acknowledgments

We thank Franck Lamberton, Danielle Ibarrola, Thomas Bastelica, Jessica Léone, Justine Epinat, and Andrea Díaz-Barriga Yáñez for their help with data collection. We are also grateful to the children and parents who participated. This study was partially supported by MEXT/JSPS KAKENHI (JP24H02172 and JP24H01559) and JST FOREST Program (JPMJFR231V) for T.N., Fondation de France (00123415 / WB-2021-38649), Fédération pour la Recherche sur le Cerveau (AP-FRC-2022), and Agence Nationale de la Recherche (ANR-23-CE28-0002-01) for J.P. The funders had no role in the study design, data collection and analysis, decision to publish, or preparation of the manuscript.

## Supplementary Information

### Prediction accuracy in other anatomical ROIs

We evaluated between-group differences in other anatomical ROIs that have been frequently found implicated in previous studies of mental arithmetic in adults and children (Istomina & Arsalidou, 2024; Sokolowski et al., 2023). These included the left inferior frontal gyrus (IFG), the hippocampus, the angular gyrus (AG), and the intraparietal sulcus (IPS). In the opercular part of the left inferior frontal gyrus (IFG), we found significantly higher prediction accuracy for adult participants than the other groups (Kruskal-Wallis test: *P* < 0.001, *η^2^* = 0.162; Wilcoxon rank-sum test: Adult vs. 14-yo, *P* < 0.001, *d* = 1.00; Adult vs. 11-yo, *P* < 0.001, *d* = 1.12; Adult vs. 8-yo, *P* = 0.006, *d* = 0.75; **Supplementary Figure S1A**). In contrast, we found a significant main effect of age and larger prediction accuracy for 11-yo than adults in the right hippocampus (Kruskal-Wallis test: *P* = 0.035, *η^2^* = 0.049; 11-yo vs. Adult, *P* = 0.005, *d* = 0.76; **Supplementary Figures S1B**), but no significant main effect was found in the left hippocampus (*P* = 0.071, *η^2^* = 0.065; **Supplementary Figures S1C**). In the right AG, we found a significant main effect of age groups, but post-hoc tests did not show a monotonic increase with age (Kruskal-Wallis test: *P* = 0.019, *η^2^* = 0.081; Wilcoxon rank-sum test: adults vs. 14-yo, *P* = 0.0049, *d* = 0.72; 11-yo vs. 14-yo, *P* = 0.0034, *d* = 0.70; **Supplementary Figures S1E**). No clear between-group difference was observed in the left AG (*P* = 0.1436, *η^2^* = 0.044; **Supplementary Figures S1D**), left IPS (*P* = 0.594, *η^2^* = 0.012; **Supplementary Figures S1F**), and right IPS (*P* = 0.237, *η^2^* = 0.075; **Supplementary Figures S1G**). Overall, these results indicate that the age-related increase in similarity with the ANN is primarily seen in left frontal areas in older participants, though a similarity between the ANN and the right hippocampus may also be seen earlier in development.

### Effect of tie problems and sum-to-10 problems

In addition to small and large problems, previous studies have reported faster reaction times for tie problems (e.g., “3+3”) and sum-to-10 problems (e.g., “3+7”) compared to other addition problems (Aiken & Williams, 1973; Groen & Parkman, 1972). Indeed, these problems may have a special status in memory (Chen & Campbell, 2018; LeFevre, Shanahan, & DeStefano, 2004). We therefore tested whether ties and sum-to-10 problems demonstrate better prediction patterns compared to other problems using the ANN model. We did not find any significant difference for these problems compared to other problems in any groups (**Figure S4A-D**). In the anatomical L. PreCS ROI, no age group showed larger prediction accuracy in ties and sum-to-10 problems than other problems (Wilcoxon sign-rank test: *P* > 0.227 for all age groups; **Figure S4E**). These results indicate that the ANN model does not show better prediction performance for activity associated with ties and sum-to-10 problems compared to other problems.

**Figure S1.**
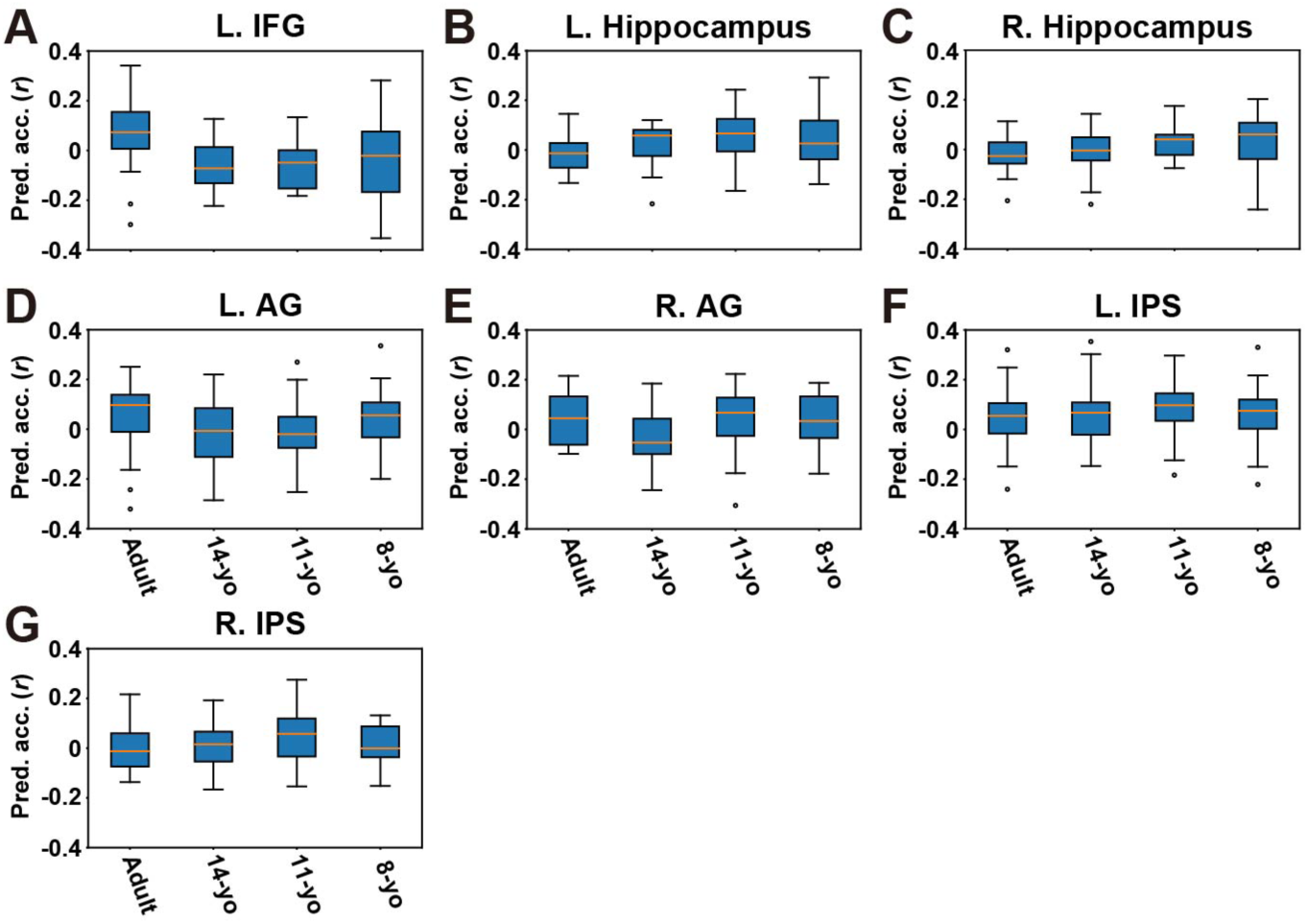
Prediction accuracy using the ANN model in anatomical regions. Box plots show the prediction accuracy of four age groups using the ANN model, averaged in the anatomical (**A**) left inferior frontal gyrus (L. IFG), (**B**) L. hippocampus, (**C**) right (R) hippocampus, (**D**) left angular gyrus (L. AG), (**E**) R. AG, (**F**) L. intraparietal sulcus (IPS), and (**G**) R. IPS regions-of-interest (ROIs). Error bar, SD.

**Figure S2.**
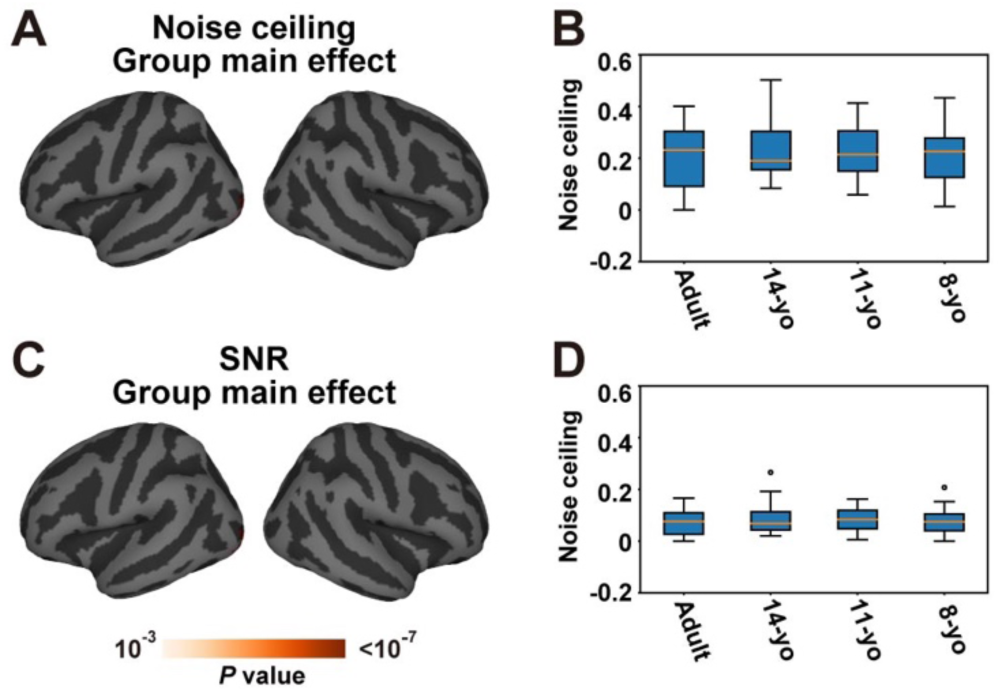
Noise analysis. Whole cortical maps of (**A**) the main effect of noise ceiling and (**C**) signal-to-noise ratio (SNR) across four age groups (Kruskal-Wallis test). Statistical significance is evaluated with a peak-level *P* < 0.001 and cluster-level *P* < 0.05 (false discovery rate [FDR] corrected). Box plots show the (**B**) noise ceiling and (**D**) SNR of four age groups using the ANN model, averaged in the anatomical left precentral sulcus (L. PreCS) ROI.

**Figure S3.**
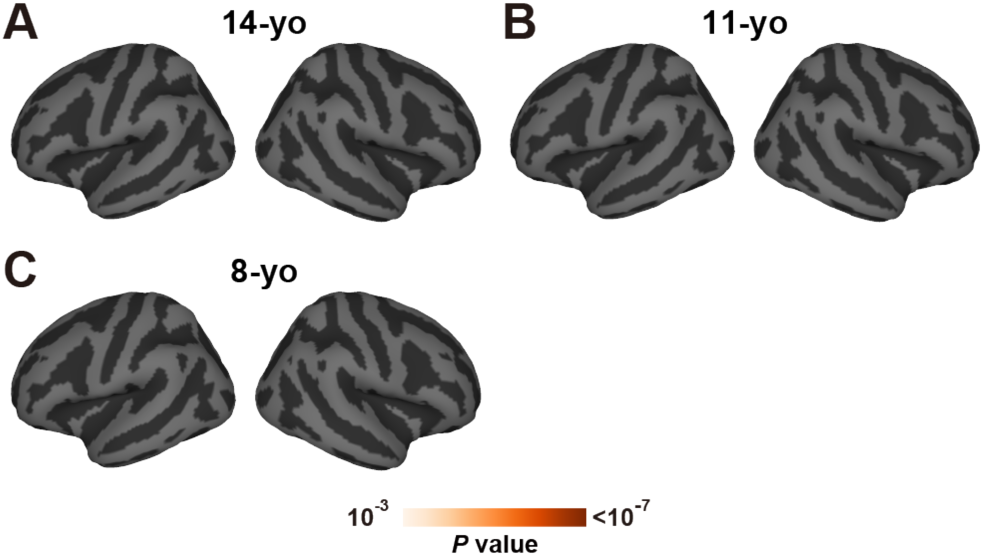
Small vs. large problems. Whole cortical maps of the direct comparison of prediction accuracy between small and large problems using the ANN model (Wilcoxon sign-rank test), separately analyzed for (**A**) 14-yo, (**B**) 11-yo, and (**C**) 8-yo. Statistical significance is evaluated with a peak-level *P* < 0.001 and cluster-level *P* < 0.05 (FDR corrected).

**Figure S4.**
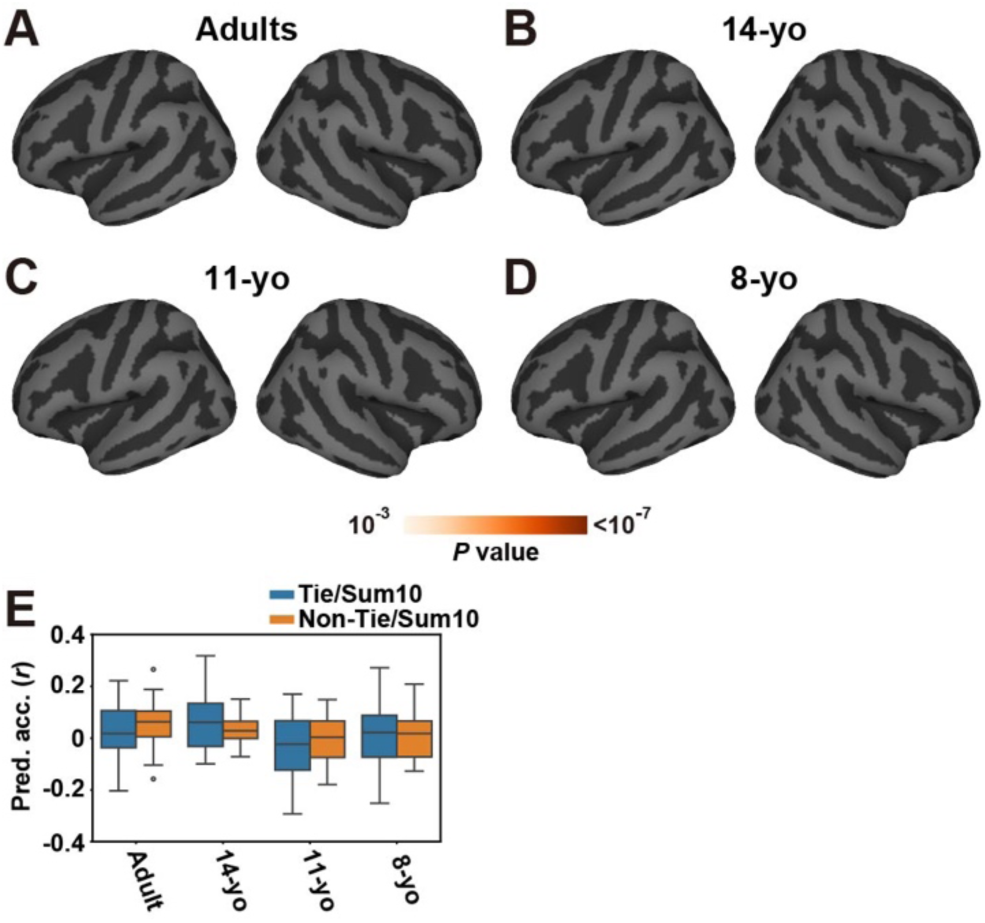
Tie/sum-to-10 vs. non-tie/sum-to-10 problems. Whole cortical maps of the direct comparison of prediction accuracy between tie/sum-to-10 and non-tie/sum-to-10 problems using the ANN model (Wilcoxon sign-rank test), separately analyzed for (**A**) adults, (**B**) 14-yo, (**C**) 11-yo, and (**D**) 8-yo. Statistical significance is evaluated with a peak-level *P* < 0.001 and cluster-level *P* < 0.05 (FDR corrected). (**E**) A box plot shows prediction accuracy of four age groups, separately calculated for tie/sum-to-10 and non-tie/sum-to-10 problems, averaged in the anatomical L. PreCS ROI.

**Figure S5.**
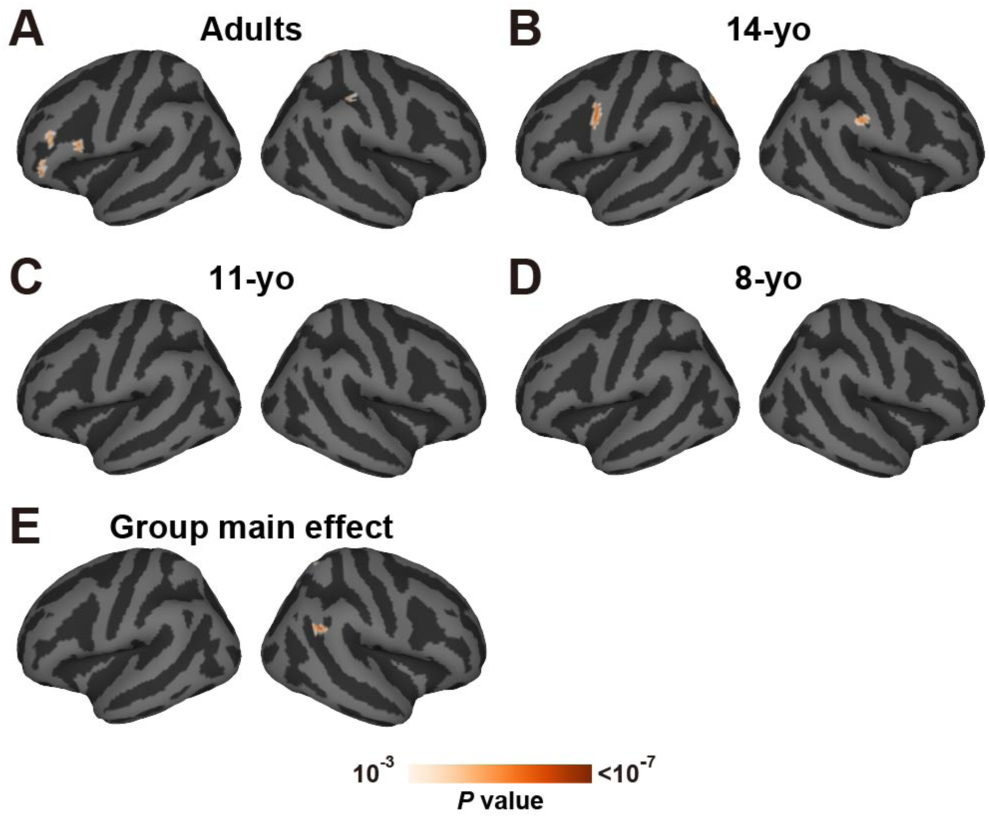
Prediction performance of the discrete model. Prediction accuracy of the discrete model, mapped onto the cortical surface for (**A**) adults, (**B**) 14-yo, (**C**) 11-yo, and (**D**) 8-yo. Only significantly predicted vertices are shown (peak-level, uncorrected *P* < 0.001; cluster-level, FDR corrected *P* < 0.05). (**E**) Whole cortical maps of the main effect of prediction accuracy across four age groups (Kruskal-Wallis test).

**Figure S6.**
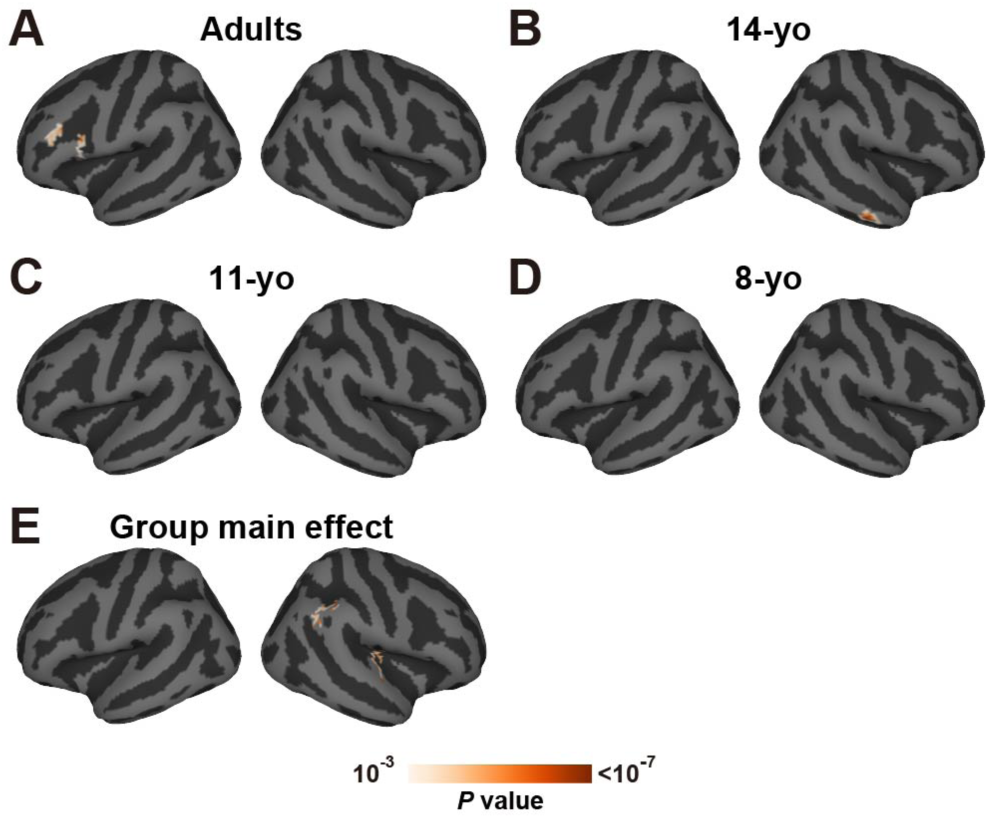
Prediction performance of the problem-size model. Prediction accuracy of the problem-size model, mapped onto the cortical surface for (**A**) adults, (**B**) 14-yo, (**C**) 11-yo, and (**D**) 8-yo. Only significantly predicted vertices are shown (peak-level, uncorrected *P* < 0.001; cluster-level, FDR corrected *P* < 0.05). (**E**) Whole cortical maps of the main effect of prediction accuracy across four age groups (Kruskal-Wallis test).

**Figure S7.**
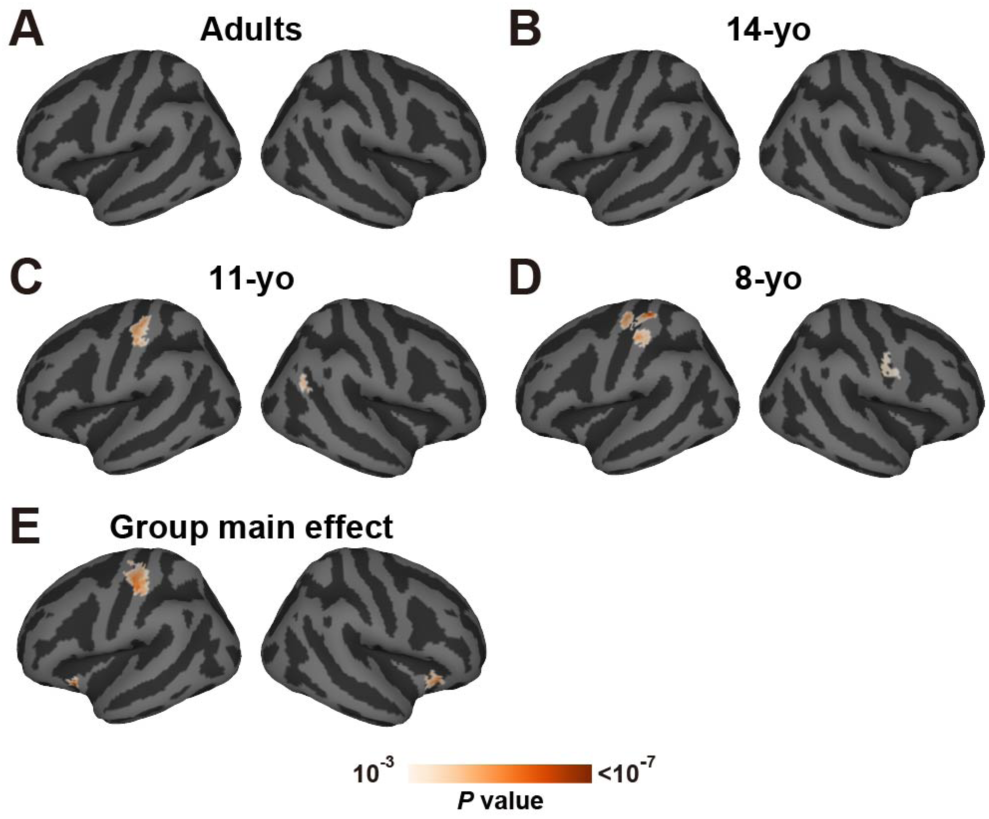
Prediction performance of the difficulty model. Prediction accuracy of the difficulty model, mapped onto the cortical fuigsurface for (**A**) adults, (**B**) 14-yo, (**C**) 11-yo, and (**D**) 8-yo. Only significantly predicted vertices are shown (peak-level, uncorrected *P* < 0.001; cluster-level, FDR corrected *P* < 0.05). (**E**) Whole cortical maps of the main effect of prediction accuracy across four age groups (Kruskal-Wallis test).

**Figure S8.**
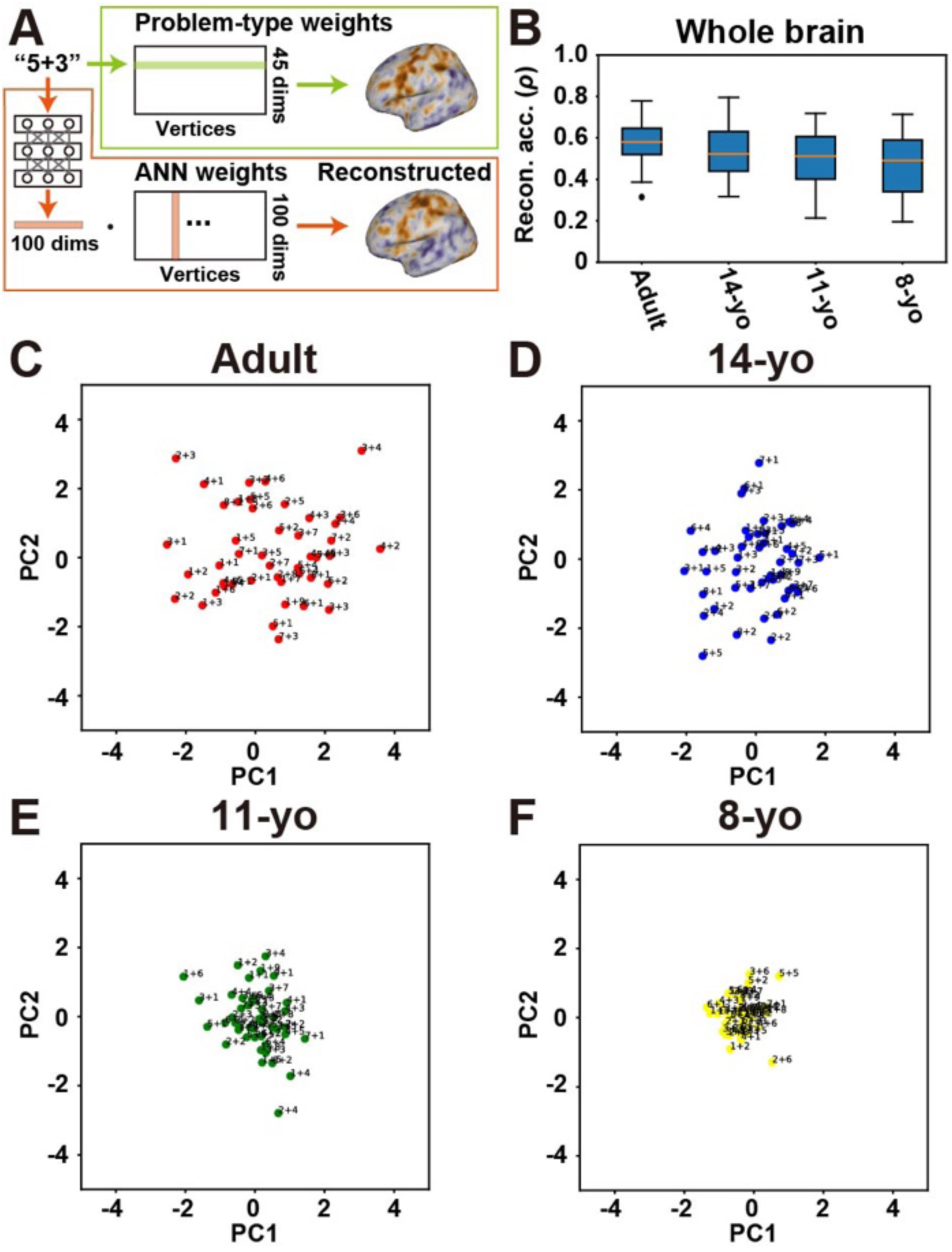
Principal component analysis using reconstructed weights. (**A**) Schematic image of weight reconstruction analysis. A weight vector corresponding to the addition problem “5+3” is extracted from the weight matrix of the discrete model. This weight vector is reconstructed by multiplying ANN features to the weight vectors of the ANN model. (**B**) The whole-cortex reconstruction accuracy is evaluated based on Spearman’s correlation coefficient between the original and reconstructed weight vectors, and box plots show average reconstruction accuracy across 45 problems (**C-F**) Principal component analysis (PCA) based on the reconstructed weights by the ANN model. 45 addition problems are mapped onto the two-dimensional space of the first and second principal components (PC1 and PC2, respectively) based on their loadings, separately visualized for (**C**) adults, (**D**) 14-yo, (**E**) 11-yo, and (**F**) 8-yo participants.

**Figure S9.**
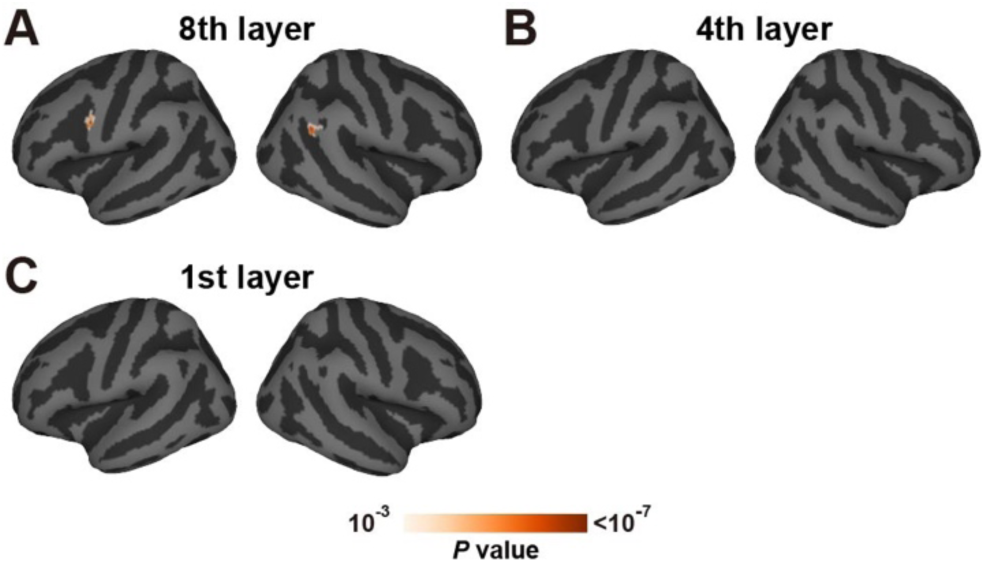
Whole cortical maps of the main effect of prediction accuracy across four age groups (Kruskal-Wallis test), separately shown for the models using (**A**) 8^th^ ANN layer, (**B**) 4^th^ ANN layer, and (**C**) 1^st^ ANN layer. Statistical significance is evaluated with a peak-level *P* < 0.001 and cluster-level *P* < 0.05 (FDR corrected).

